# Modulatory Effects of M3 Muscarinic Acetylcholine Receptor on Inflammatory Profiles of Human Memory T Helper Cells

**DOI:** 10.1101/2024.09.08.611875

**Authors:** Fatemeh Gholizadeh, Mehri Hajiaghayi, Jennifer S. Choi, Samuel R. Little, Niloufar Rahbari, Kelly Brotto, Eric Han, Steve C.C. Shih, Peter J. Darlington

**Affiliations:** Department of Biology, Concordia University, Montréal, Québec, Canada; Department of Medicine, McGill University, Montréal, Québec, Canada; Department of Electrical and Computer Engineering, Center of applied synthetic biology, Concordia University, Montréal, Québec, Canada; Department of Health, Kinesiology & Applied Physiology, Concordia University, Montréal, Québec, Canada

**Keywords:** Memory T helper Cells, Oxotremorine-M, IFN-γ, IL-17A, IL-4, M3 Muscarinic Acetylcholine Receptor, NF-κB p65

## Abstract

Memory T helper (Th) cells, generated after immunogenic challenge, are crucial in directing the adaptive immune response. Muscarinic ACh receptor (mAChR) subtypes expressed by immune cells can be stimulated with acetylcholine or muscarinic-selective drug oxotremorine-M. Cholinergic signaling can influence immune cells, but it is not known how cholinergic stimuli regulate memory Th cells. This study focused on the role of mAChRs, specifically the M3 muscarinic ACh receptor (M3R), in the cytokine profile and NF-κB p65 activity of primary human memory Th cells.

Memory Th cells (CD3^+^CD4^+^CD45RA^-^CD45RO^+^) were isolated from healthy participants’ peripheral blood. Cell culture was performed with anti-CD3/anti-CD28/anti-CD2 reagent, oxotremorine-M (M1R-M5R agonist), atropine (M1R-M5R antagonist), and J104129 (M3R-selective antagonist). MR1-MR5 genes *CHRM1*-*CHRM5* were measured with RT-qPCR. Protein expression of M3R and phosphorylated NF-κB p65 were quantified by Western blot. The secretion of IFN-γ, IL-17A, and IL-4 was assessed by ELISA and intracellular cytokine staining flow cytometry. *CHRM3*, encoding M3R, was knocked out using CRISPR-Cas9 gene targeting.

Memory Th cells expressed all five mAChR subtypes. Oxotremorine-M increased IFN-γ and IL-17A while reducing IL-4 in an atropine-sensitive manner. Stimulation of mAChRs in cells with *CHRM3*-knockout or M3R blockade prevented increases in IFN-γ and IL-17A but continued to inhibit IL-4. mAChR stimulation enhanced NF-κB p65 activity without affecting cell proliferation, viability, or M3R expression.

This investigation demonstrates that muscarinic signaling increases the pro-inflammatory profile of memory Th cells, including NF-κB p65, IFN-γ, and IL-17A, with a reduction in IL-4. Focusing on M3R blockers could modulate adaptive immune responses and alleviate immune-related conditions.

## Introduction

Acetylcholine (ACh) is primarily known for its role as a neurotransmitter in the peripheral parasympathetic (cholinergic) branch of the autonomic nervous system. It binds to a family of muscarinic receptors (mAChRs) comprised of five subtypes: M1R, M2R, M3R, M4R and M5R encoded by the genes *CHRM1, CHRM2, CHRM3, CHRM4,* and *CHRM5*, respectively. Although these receptors are present in neuronal tissues, they are also expressed by non-neuronal tissues including immune cells (1–3). The M1R, M3R, and M5R subtypes preferentially couple to G_q/11_ proteins, leading to activation of phospholipase Cβ, resulting in mobilizing Ca^2+^ from intracellular stores and activating protein kinase C (PKC). M2R and M4R couple to G_i/o_ proteins, leading to inhibition of adenylyl cyclase and thus reduced cAMP formation (4).

During adaptive immunity, a fraction of effector T helper (Th) cells transition into memory Th cells, primed to mount rapid and robust secondary immune responses upon re-exposure. Th cells exhibit various profiles such as Th1, Th2, Th17, Th22, and Tfh, each characterized by specific cytokine secretion patterns and effector functions. Elevated levels of Th1 cells producing IFN-γ and Th17 cells producing IL-17A are consistently observed across a spectrum of autoimmune diseases considered to be pro-inflammatory (5–7). Th2 cells, which express IL-4, play an anti-inflammatory role. Autoimmune diseases are often characterized by reduced IL-4 production, prompting therapeutic strategies aimed at reinstating a balanced ratio of Th1/Th17 to Th2 cytokines to physiological levels (8, 9). Activation of Th cells occurs in part via downstream phosphorylation of nuclear factor κB (NF-κB) composed of p65/p50 dimers (10, 11). NF-κB p65 signalling is required for immune cell function through transcriptional regulation of cytokine expression such as IL-17A and IFN-γ (12, 13).

Th cells express mAChRs and other cholinergic components, including nicotinic receptors, acetylcholinesterase, and choline acetyltransferase (14–16). Choline acetyltransferase is expressed by Th17 cells and primes them for enhanced autoimmunity in experimental autoimmune encephalomyelitis (17). Oxotremorine-M is an mAChR agonist of all M1R-M5R, atropine is an antagonist for M1R-M5R (18), and J104129 is a selective M3R antagonist (19). These pharmacological agents are primarily utilized in research to study the cholinergic nervous system and to elucidate the impact of acetylcholine on immune cell functions. ACh induced Ca^2+^ signaling and increased the expression of c-fos leading to the release of IL-6 in mouse spleen cells (20). In human T cells, ACh induced the release of IL-2, as well as an increase in IL-2 receptor expression (21). M3R was required for optimal adaptive immune responses and immunological memory following helminth and bacterial infections in a mouse model (22). M3R is implicated in the production of TNF-α in the mouse alveolar macrophage through their ability to induce the degradation of IκBα, which indicated NF-κB activation (23). Moreover, a recent article showed that cholinergic markers coincided with severe COVID-19 infections and that glucocorticoid treatment reduced ACh levels (24). While the pro-inflammatory role of cholinergic signaling in Th cells is acknowledged, its precise contribution to memory Th cell modulation is incompletely understood

This study investigated the expression and function of muscarinic receptors in memory Th cells exhibiting Th1, Th2 and Th17 profiles. Stimulation of mAChRs increased IFN-γ and IL-17A cytokines in an M3R-dependent manner with concurrent NF-κB p65 activation, while it inhibited anti-inflammatory IL-4 cytokine in an atropine-sensitive, M3R-independent manner. Thus, muscarinic receptor stimulation favors a pro-inflammatory profile in memory Th cells.

## Material and methods

### Peripheral blood lymphocyte preparation

Healthy participants were determined through self-reporting during semi-structured interviews. Exclusion criteria comprised individuals under the age of 18, those diagnosed with medical conditions, individuals currently taking adrenergic, cholinergic, or steroid-based medications. Participants were rescheduled if they reported vaccinations or used recreational drugs within 2 weeks prior to the blood draw. Participants provided informed signed consent prior to the study and filled a basic demographic questionnaire. Venous blood was drawn by a phlebotomist with a 21-gauge butterfly needle in antecubital fossa, collected into heparinized green top blood collection vials (Fisher Scientific, Ottawa, Canada). Peripheral blood mononuclear cells (PBMCs) obtained from participants using 1.077-1.080 g/m density lymphocyte separation media (Wisent Bioproducts, st-Bruno, Canada) according to a published protocol (25). PBMCs were cryopreserved at −80^◦^C to be used for purification of memory Th cells.

### Enrichment and activation of memory Th cells

Memory Th cells were enriched from PBMC samples by immunopanning negative selection with EasySep™ human memory CD4^+^ T cell enrichment kit (Stemcell Technologies, Vancouver, Canada) following the protocol provided by manufacturer. Briefly, cells expressing CD8, CD14, CD16, CD19, CD20, CD36, CD45RA, CD56, CD123, TCRγ/δ, CD235a (glycophorin A) were depleted using the enrichment kit and EasySep™ magnet. Isolated memory Th cells purity was determined by flow cytometry staining for CD3, CD4, CD45RA and CD45RO. Memory Th cells were cultured in RPMI 1640 supplemented with 10% fetal bovine serum (FBS), 2 mM L-glutamine, 100 U/mL penicillin, and 100 μg/mL streptomycin (Wisent Bioproducts) in 5% CO_2_ humidified incubator at 37^◦^C. Memory Th cells were plated at 1.5 × 10^6^ cells/mL in 96-well U-bottomed tissue plates (Fisher Scientific). Cells stimulated with 3.125 µL/mL ImmunoCult reagent which is an antibody complex of anti-CD3, anti-CD28, and anti-CD2 (Stemcell Technologies). Cells were cultured with atropine at 10 µM (Millipore Sigma Aldrich, Oakville Canada) for 30 mins and then were exposed to oxotremorine-M at 100 µM (Cayman Chemicals from Cedarlane, Burlington Canada) for an additional 5 days. M3 antagonist J104129 at 0.01 µM (Tocris Bioscience, from Cedarlane) was added 30 mins in advance, then oxotremorine-M (100 µM) for 5 days. Cell free supernatants were collected, centrifuged at 493g 7 mins room temperature, and used for analysis with ELISA.

### Enzyme-Linked Immunosorbent Assay (ELISA)

IFN-γ, IL-4 (BD Biosciences, USA), and IL-17A were measured from memory Th cell supernatants using commercially available sandwich ELISA kits according to manufacturer protocols (ThermoFisher Scientific, Burlington, Canada). Modifications included 75µL of coating and capture antibody per well, and blocking with ELISA blocking buffer powder (Bethyl Laboratories Inc. from Cedarlane) reconstituted in milliQ H_2_O. TMB substrate for the IL-4 and IFN-γ was ELISA Ultrasensitive TMB Substrate for HRP (Applied Biological Mat.Inc. Products from Cedarlane). Absorbances at 450nm and correction with 570nm were quantified with a microplate reader (Bio Tek, BD Bioscience, Mississauga Canada).

### Flow cytometry

Antibodies were anti-CD3-PerCP (clone HIT3a, Biolegend, San-Diego USA), anti-CD4-APC (clone RPA-T4, BD Biosciences), anti-CD45RO-PE (clone UCHL1, ThermoFisher Scientific), anti-CD45RA-FITC (clone HI100, ThermoFisher Scientific), anti-IL-17A-PE (clone BL168, eLabscience from Cedarlane), anti–IFN-γ- BV421 (clone B27, BD Biosciences), anti-IL-4-PE-Cy7 (clone 8D4-8, ThermoFisher Scientific), PE mouse IgG1 κ isotype (clone MOPC-21, BD Biosciences), BV421 mouse IgG1 κ isotype (clone X40 BD Biosciences), and PE-Cy7 mouse IgG1 κ isotype (clone P3.6.2.8.1, ThermoFisher Scientific). The flow cytometer was FACS Verse system (BD Biosciences). For intracellular cytokine staining, after 5 days of incubation, memory Th cells were re-stimulated with a mixture of phorbol 12 myristate 13 acetate (PMA) at 50 ng/mL, ionomycin at 500 ng/mL, and brefeldin A at 10 µg/ml (Millipore Sigma Aldrich) for 5 h at 37°C. Cells were centrifuged (493 g, 7 mins, room temperature) and the supernatant was removed. Cells were stained for CD4 for 30 mins on ice, fixed and permeabilized with Cytofix/Cytoperm kit (BD Biosciences), then stained with anti-IFN-γ, anti-IL-17A, or anti-IL-4. Isotype matched control antibodies enabled the correct background and confirmed antibody specificity. Compensation was determined with single stained samples prior to the analysis. To determine the percentage of cells producing selected cytokines, values obtained with isotype controls were subtracted from those with specific mAb. All samples were analyzed using FlowJo software (BD Bioscience, USA).

### Proliferation with CFDASE labelling, viability with 7AAD

For proliferation, human memory Th cells were labelled with 0.05 mM carboxyfluorescein diacetyl succinimidyl ester (CFDASE) cell proliferation reagent (Millipore Sigma Aldrich) according to published protocol (26). PBS with 5% FBS stopped the reaction, and washed three times. Cells at 1.5 ×10^6^ cells/mL were cultured for 5 days under similar experimental conditions as previously described, including the presence of oxotremorine-M (100 µM), with or without 10 µM atropine. Post-incubation, the cells underwent a 30-mins CD4 staining process on ice and were then analyzed using flow cytometry. Viability was evaluated via using 7 aminoactinomycin D, 7AAD-PerCP at 1:100 in PBS (ThermoFisher Scientific) to stain the dead cells.

### Quantitative RT-PCR

Total RNA was extracted from 2 ×10^6^ memory Th cells using PureLink™ RNA Mini Kit (Invitrogen™, Carlsbad, USA) according to the manufacturer’s protocols. Isolated RNA was quantified using a spectrophotometer (NanoDrop™ 2000c, ThermoScientific). RNA samples had A260/A280 ratio of ∼2.0. Next, 1400 ng of total RNA was reverse transcribed into 40 µL complementary DNA using iScript™ Reverse Transcription Supermix for RT qPCR (Bio Rad Laboratories, Hercules, USA). Reaction was in programmable Thermal Controller (Bio-Rad Laboratories st-Laurent, Canada). Each complementary DNA sample was diluted to 7 µg/mL and were amplified using the TaqMan™Fast Advanced Master Mix (Applied Biosystems™, Foster City, USA) according to the manufacturer’s protocol. TaqMan™ gene probes were used to measure mAChR subtypes including *CHRM1* (Hs00265195_s1), *CHRM2* (Hs00265208_s1), *CHRM3* (Hs00265216_s1), *CHRM4* (Hs00265219_s1), and *CHRM5* (Hs00255278_s1). Peptidylprolyl isomerase A (PPIA, Hs99999904_m1) was used as an internal standard. 3 µL of complementary DNA was used per 20 µL of amplification reaction. Assay was performed in triplicate. Thermal cycling had activation step of 50^◦^C for 2 mins and 95^◦^C for 20 sec, followed by 50 cycles of 95^◦^C for 3 sec and 60^◦^C for 30 sec. (CFX 96 Real Time System C1000 Thermal Cycler Bio-Rad Laboratories). Data were normalized to reference gene PPIA. In order to determine the relative expression of all the muscarinic receptors, the muscarinic receptor data was normalized to 1 for *CHRM1* using 2 ^-(ΔΔCt)^ method.

### CRISPR-Cas9 ribonucleoprotein (RNP) knockout of CHRM3 in memory Th cells

Memory Th cells were incubated in a complete culture medium with 100 IU/mL recombinant human IL-2 (Fisher Scientific, Ottawa, ON) for 24 hrs in a 5% CO_2_ at 37°C, then activated with human T-Activator CD3/CD28 Dynabeads (Fisher Scientific, Ottawa, ON, #11131D) for 48 hrs. After bead removal and a 24-hr rest, cells were resuspended in Neon electroporation buffer. Memory Th cells were resuspended in the Neon electroporation buffer (ThermoFisher Scientific). Gene targeting was with Cas9 2NLS nuclease, single guide (sg) RNAs for T cell receptor alpha constant (*TRAC*) as the positive control, *CHRM3*, and a non-targeting sgRNA (Synthego, Redwood City USA). The payload sequences for the multi sgRNAs were as follows: *TRAC*: 5’-CUCUCAGCUGGUACACGGCA-3’, 5’-GAGAAUCAAAAUCGGUGAAU-3’, and 5’- ACAAAACUGUGCUAGACAUG-3’; *CHRM3*: 5’-GGUUGUACUGUUAUUGUGCA-3’, 5’- UCCGAUGCAGGGCUGCCCCC-3’, and 5’-AUCGGUGGUACCGUCUGGAG-3’.

The protocol for CRISPR knockout experiments on the human primary T cells was according to published methods (27–29). Briefly, 100 pmol of sgRNA and 50 pmol of Cas9 were mixed per 1 × 10^6^ memory Th cells and incubated for 10 mins to form Cas9 RNP complexes. Memory Th cells were mixed with Cas9 RNP complexes containing either scrambled sgRNA or multi-sgRNA targeting the *CHRM3* or *TRAC* genes in the electroporation buffer, resulting in 2 × 10^7^ cells/mL, and electroporated using Neon transfection system (ThermoFisher Scientific). 100 μL of the cells and Cas9 RNP mixture containing 2 × 10^6^ cells was transferred to the Neon capillary electroporation tip and electroporated using the manufacturer’s recommended parameters for Th cells (1600 V, 10 ms, 3 pulses). Cells were placed in pre-warmed culture media with 400 IU/mL recombinant human IL-2 for recovery. Knockout efficiency was determined after 4 days using fluorescence-based cell counting for *TRAC* calculated by dividing the number of living cells with low fluorescence by the total number of living cells, and multiplying by 100. RT-qPCR data analysis for *CHRM3* calculated by comparing the relative expression levels of the target gene in knockout cells to control cells and multiplying by 100. For cell culture, 0.3 × 10^6^ memory Th cells were plated per well in 96-well U-bottomed tissue culture plates. The cells were left unstimulated or stimulated with 3.125 µL/mL ImmunoCult reagent and/or oxotremorine-M at 100 µM for 5 days. Cell-free supernatants were collected for ELISA.

### Primary T cell lysis and western blot

Memory Th cell blast was made by incubating for 24 hrs with 1 ng/mL PMA, 1 ng/mL ionomycin, followed by washing and 24 hrs rest in incubator. The cells were incubated at 1.5 x 10^6^ cells/mL with no activation, or activation with 3.125 µL/mL ImmunoCult. Oxotremorine-M was added at 100 µM. The cells were lysed after 15 mins, with triton X buffer (150 mM NaCl, 10 mM Tris pH 7.5, 1% triton X-100, 1 mM Na_3_VO_4_, 5 mM EDTA, 10 μg/mL leupeptin, 10 μg/mL aprotinin, 25 μM p-nitrophenyl p’-guanidinobenzoate, 1X protease inhibitor cocktail (Roche, Laval, Canada). The debris was pelleted for 5 mins at 2500 g in 4^◦^C on ice. 2X Laemmli buffer with 5% β-mercaptoethanol was added to the supernatant and boiled in 95 ^◦^C for 5 mins. 20 µg of total protein, quantified by detergent compatible protein assay kit II (Bio-Rad Laboratories), and microplate reader (Bio Tek, BD Bioscience) were separated by 10% SDS-PAGE electro transferred to nitrocellulose membranes (BioRad Laboratories). Membranes were blocked at 4 ^◦^C overnight in Tris-buffered saline (20 mM Tris-HCl, 150 mM NaCl, pH 7.6) containing 0.1% Tween-20 (TBS-T), supplemented with 5% non-fat milk (Anatol Spices, Montreal Canada). Membranes were washed in TBS-T and incubated overnight at 4 ^◦^C with monoclonal pNF-κB p65 (Ser 529)-specific antibody (1:500 dilution, clone 817403, Biolegend), monoclonal NF-κB p65-specific antibody (1:2500 dilution, clone 465003, Biolegend), polyclonal CHRM3-specific antibody (1:2000 dilution, rabbit polyclonal PA5-72320, ThermoFisher Scientific), or monoclonal GAPDH antibody (1:5000 dilution, clone 10B4E3, Cusabio) in 5% milk/TBS-T. After washing with TBS-T, membranes were incubated with a secondary goat anti-mouse IgG HRP (1:3000 dilution) or goat anti-rabbit IgG (1:4000 dilution) in 5% milk/TBS-T for 2 hrs at room temperature. Chemi-luminescence detection was with Clarity^TM^ western ECL Substrate and analyzed with gel imager chemidoc XRS + system the Image Lab 5.1 software application (Bio-Rad).

### Data analysis

Data are presented as the mean ± SD. One-way ANOVA or two-way ANOVA with p < 0.05 was considered as statistically significant. Tukey’s multiple comparison’s test was used to compare differences between conditions. Statistical analyses with Prism version 5.0 (GraphPad Prism, San Diego, USA).

## Results

### mRNA expression and function of mAChRs in memory Th cells

Memory Th cells were enriched from PBMCs of human participants using immuno-panning methods that yielded greater than 97% cells with the memory phenotype based on CD3^+^CD4^+^CD45RA^-^CD45RO^+^ (Figure 1 A, B). The expression of genes for the individual muscarinic receptor subtypes (*CHRM1-CHRM5*) was measured by RT-qPCR. All five muscarinic receptor subtypes were detected in the memory Th samples. In particular, *CHRM2, CHRM4*, and *CHRM5* were the most abundant in comparison to *CHRM1* and *CHRM3* (Figure 1 C). Oxotremorine-M significantly increased IFN-γ and IL-17A levels in activated memory Th cells, effects that were blocked by atropine (Figure 1 D, E). Additionally, Oxotremorine-M decreased IL-4 levels in memory Th cells, an effect also inhibited by atropine (Figure 1 F). These data demonstrated that oxotremorine-M augments IFN-γ and IL-17A, while it inhibits IL-4 from memory Th cell samples in an atropine-sensitive manner. To confirm these observations were occurring within Th cells, intracellular cytokine staining was used to detect the proportion of cells. Stimulation of memory Th cells with oxotremorine-M led to an increase in the proportion of CD4^+^IFN-γ^+^ cells (Figure 2 A-F), an increase in CD4^+^IL-17A^+^ cells (Figure 2 G-L), and a decrease in CD4^+^IL-4^+^ cells (Figure 2 M-R). The effects of oxotremorine-M on CD4^+^IFN-γ^+^ and CD4^+^IL-17A^+^ cells were blocked by atropine except CD4^+^IL-4^+^ cells which were not significantly blocked although a trend was noticed. The activated memory Th cells displayed 4–5 cycles of cell division, and oxotremorine-M did not change the division index or proliferation index compared to the activated condition (Figure 3 A-E). In memory Th cells, the reduction in IL-4 cytokine after exposure to oxotremorine-M suggested that the cellular viability could be compromised. Oxotremorine-M and atropine did not have any cytotoxic effects compared with non-activated and activated conditions on the memory Th cells (Figure 3 F-J). Thus, the fluctuations in cytokine levels caused by oxotremorine-M were not related to cellular proliferation or cytotoxicity.

**Figure 1.**
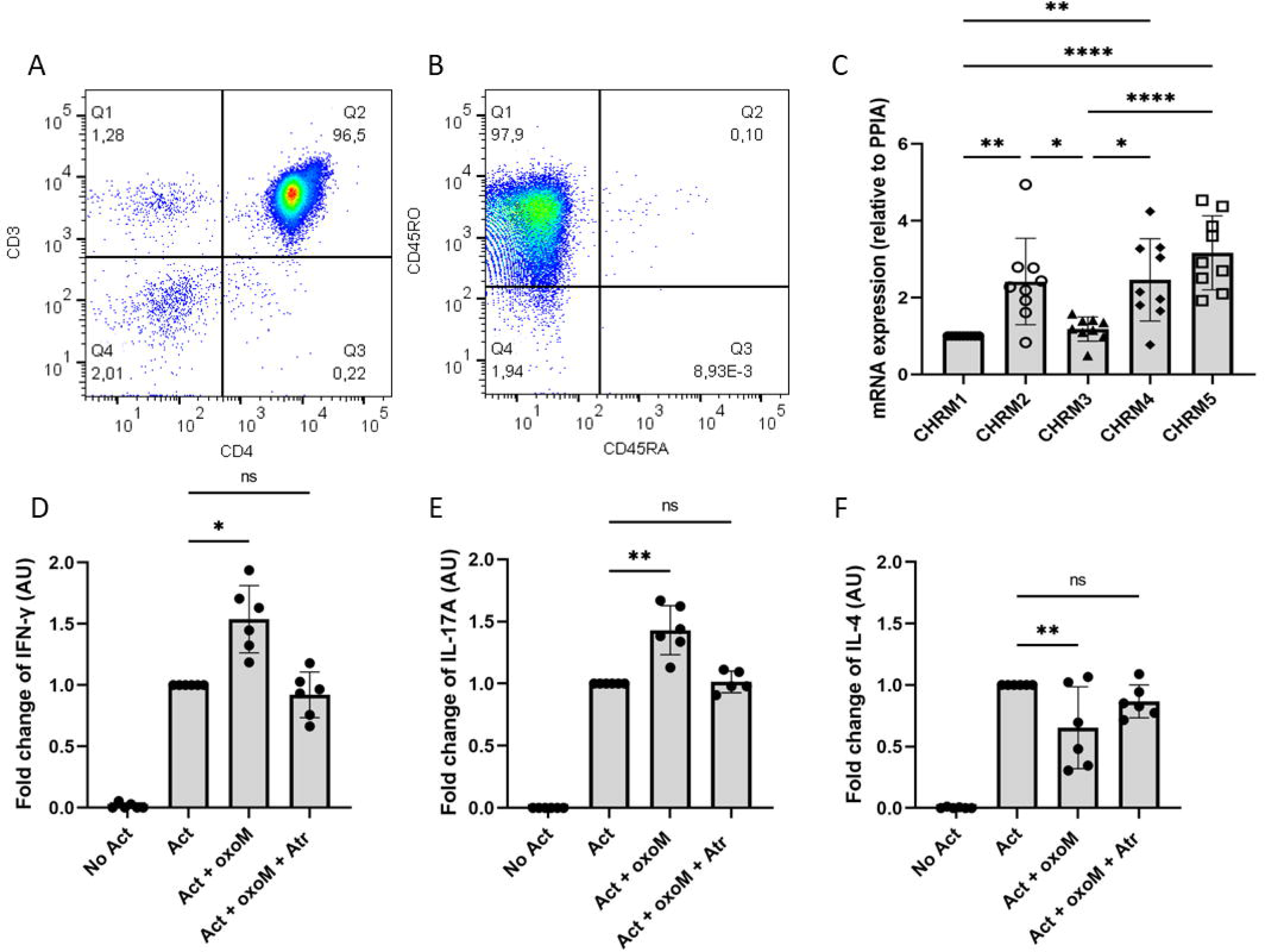
mRNA expression of M1R-M5R and the effect of muscarinic receptor stimulation on secretion of IFN-γ, IL-17A, and IL-4 from memory Th cells. (A) Representative memory Th cell expression of CD3 and CD4 measured by flow cytometry. (B) CD45RA and CD45RO plotted from quadrant Q2 from CD3^+^CD4^+^ panel. (C) mRNAs for all five muscarinic receptor subtypes were detected via RT-qPCR. Data are normalized against reference gene PPIA. The muscarinic receptor data was normalized to *CHRM1*. Aggregated data normalized to 1 for *CHRM1* is shown from memory Th cells of 9 participants. The cytokine level of (D) IFN-γ, (E) IL-17A, and (F) IL-4 measured by using ELISA. Memory Th cells were not activated (No Act); or activated with ImmunoCult reagent (Act) and incubated with oxotremorine-M or incubated with oxotremorine-M plus atropine. The data was aggregated from 6 participants. One-way ANOVA with Tukey’s multiple comparison test. *p< 0.05, **p<0.01, ****p<0.0001. ns, not significant.

**Figure 2.**
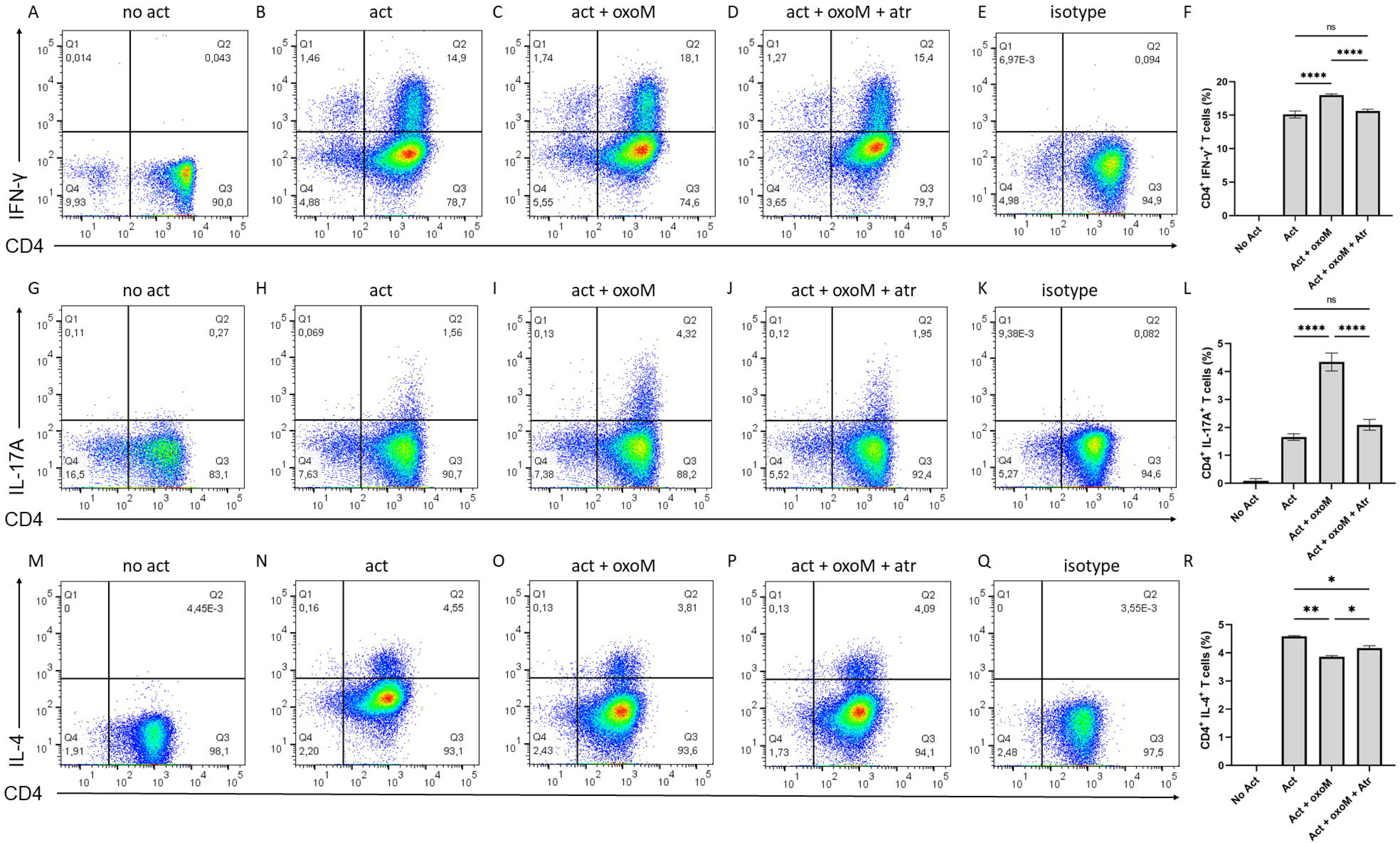
The effect of muscarinic receptor stimulation on the level of intracellular cytokines from memory Th cells. Cytokines included (A-F) IFN-γ, (G-L) IL-17A, and (M-R) IL-4. Memory Th cells were not activated (No Act); or activated with ImmunoCult reagent (Act) and incubated with oxotremorine-M or with oxotremorine-M plus atropine. An isotype control used to set gates is shown (E, K, Q). Graphs show quadrant Q2 data (CD4^+^cytokine^+^ cells) pooled together from 5 participants. One-way ANOVA with Tukey’s multiple comparison test. *p<0.05, **p<0.001, ****p<0.0001. ns, not significant.

**Figure 3.**
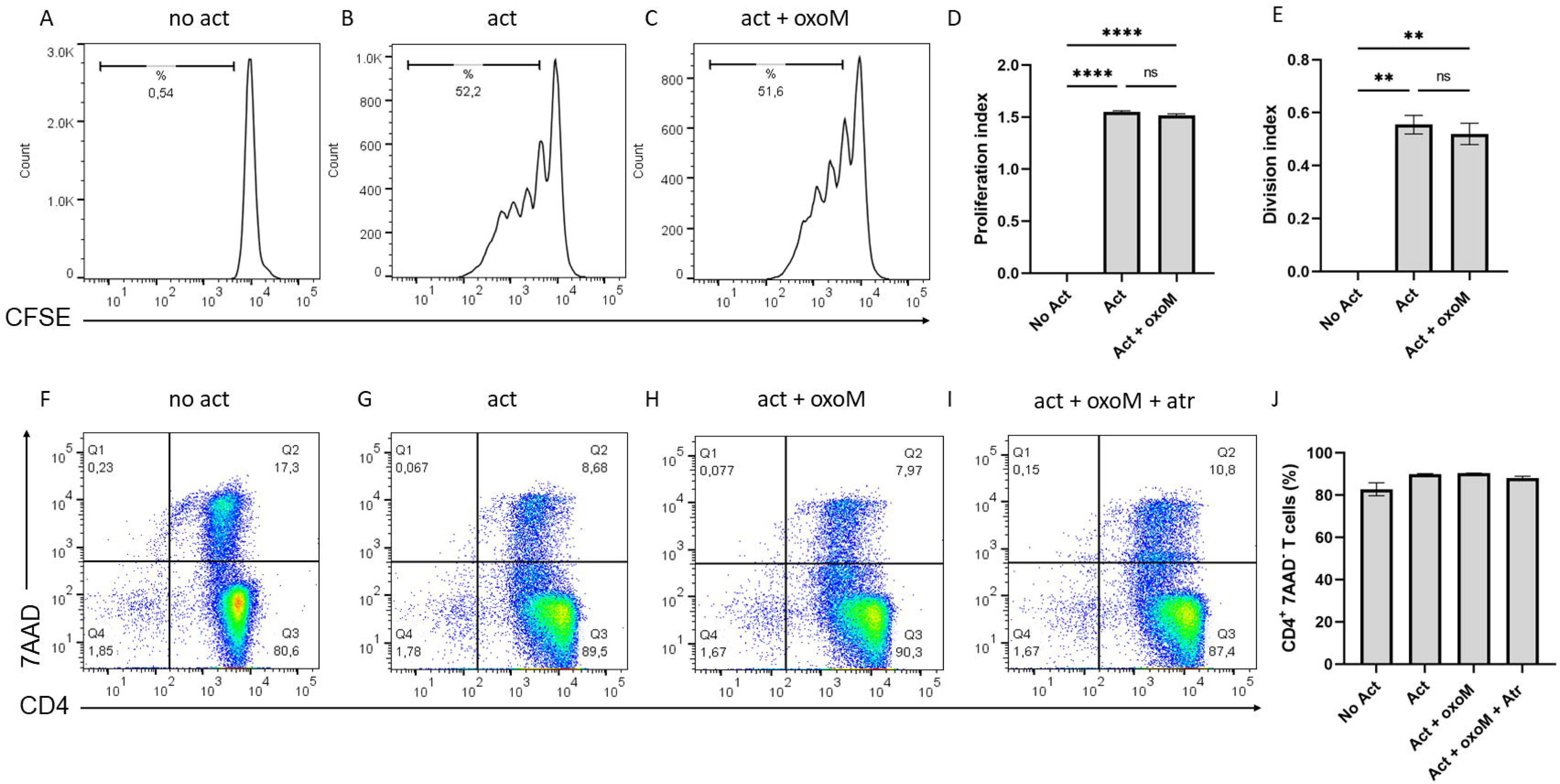
The effect of muscarinic receptor stimulation on the proliferation and viability of memory Th cells. Memory T cells were labelled with proliferation tracking dye CFDASE which diminishes with each round of cell division. Memory Th cells were (A) not activated (No Act) or (B) activated with ImmunoCult reagent (Act) and (C) incubated with oxotremorine-M. (D) Bar graphs showing division index (number of cell divisions/total cell number) and (E) proliferation index (number of cell divisions/number of divided cells). (F) Non-activated cells, (G) activated with ImmunoCult reagent, (H) activated and incubated with oxotremorine-M, and (I) activated and incubated with oxotremorine-M plus atropine. (J) Quantification of CD4^+^7AAD^-^ T cells percentage. A representative experiment from 6 participants is shown. One-way ANOVA with Tukey’s multiple comparison test. **p<0.01, ****p<0.0001. ns, not significant.

### Effects of stimulation of M3R on IFN-γ, IL-17A, and IL-4 secretion by memory Th cells

The RT-qPCR results indicated that all five muscarinic receptors were expressed, and a broad-muscarinic receptor inhibitor, atropine, blocked the pro-inflammatory effects of oxotremorine-M. To determine the contribution of M3R, a selective antagonist of M3R, J104129 (J10) was included in cell cultures. J10 effectively inhibited the oxotremorine-M-induced enhancement of IFN-γ and IL-17A (Figure 4 A-B). Although oxotremorine-M reduced IL-4, J10 failed to block the effect, leaving IL-4 levels below the activated condition (Figure 4 C). This observation reaffirmed the consistent role of M3R in governing cytokine regulation for IFN-γ and IL-17A. For IL-4, oxotremorine-M was blocked by atropine, but appeared to use a non-M3R based on the J10 experiments.

**Figure 4.**
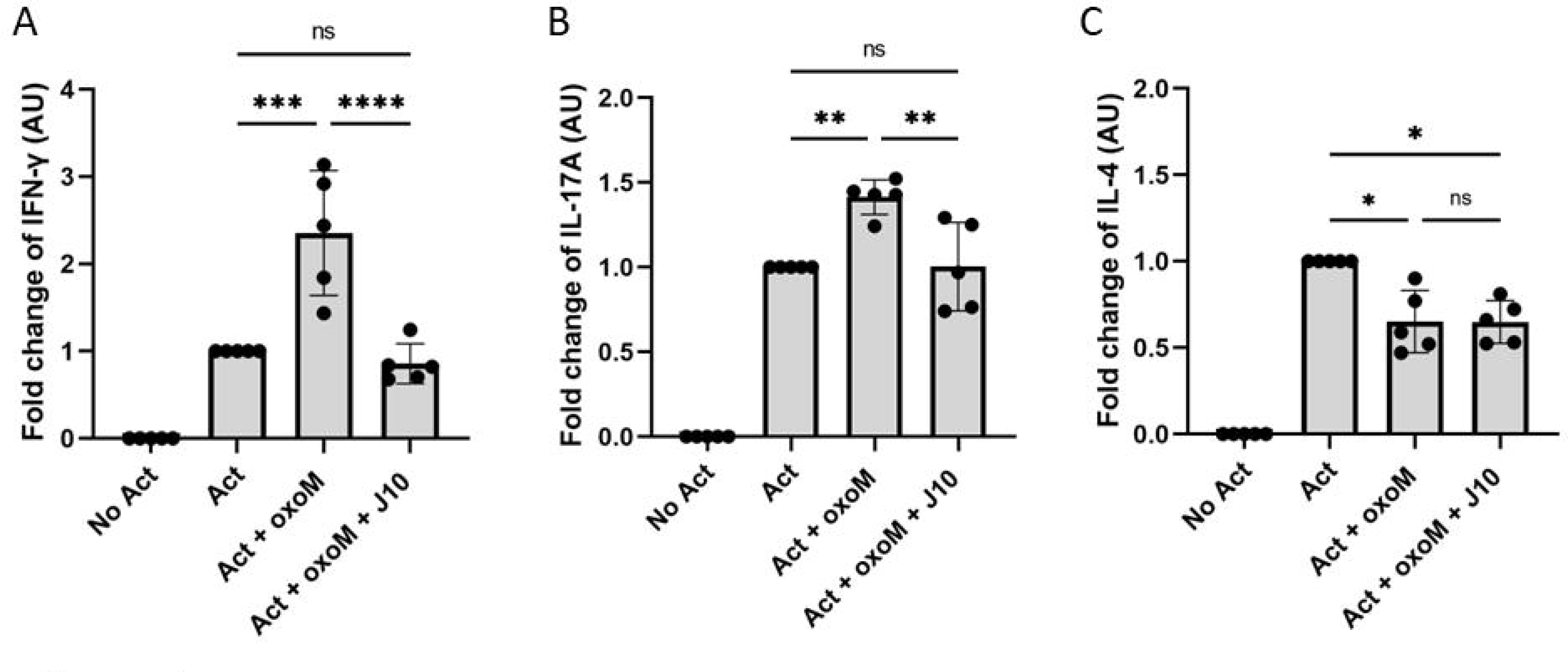
The effect of blocking M3R on secretion of IFN-γ, IL-17A, and IL-4 from memory Th cells. Memory Th cells were not activated (No Act) or activated with ImmunoCult reagent (Act) and incubated with oxotremorine-M or with oxotremorine-M plus J104129. The cytokine level of (A) IFN-γ, (B) IL-17A, and (C) IL-4 measured by ELISA. Data was aggregated from 5 participants. One-way ANOVA with Tukey’s multiple comparison test. *p < 0.05, **p<0.01, ***p<0.001, ****p<0.0001. ns, not significant.

### Effects of stimulation of mAChRs on M3R expression and phosphorylation of NF-κB p65 in memory Th cells

To assess alterations in M3R expression in activated memory Th cells following cell culture with oxotremorine-M, the levels of M3R protein and mRNA were quantified using western blot and RT-qPCR, respectively. Activation of memory Th cells downregulated M3R expression compared to the non-activated condition (Figure 5 A, B). Notably, there were no significant differences observed between the oxotremorine-M-treated and activated conditions. This finding was in line with RT-qPCR analyses demonstrating a significant reduction in M3R expression upon activation that was unaffected by oxotremorine-M (Figure 5 C). Thus, M3R was reduced during the culture period but remained detectable at day 5 of the experiment. Both IFN-γ and IL-17A cytokines are regulated by several transcription factors including NF-κB p65. To investigate whether the stimulation of mAChRs could influence NF-κB p65 activation, the phosphorylation status of serine 529 on the p65 subunit was assessed. Memory Th cells activated in the presence of oxotremorine-M displayed an increase in phosphorylation of NF-κB p65 compared to both non-activated and activated control conditions (Figure 6 A, B). Thus, muscarinic stimulation induces a pro-inflammatory profile in memory Th cells including elevated IFN-γ, IL-17A, NF-κB p65 phosphorylation and decreased IL-4.

**Figure 5.**
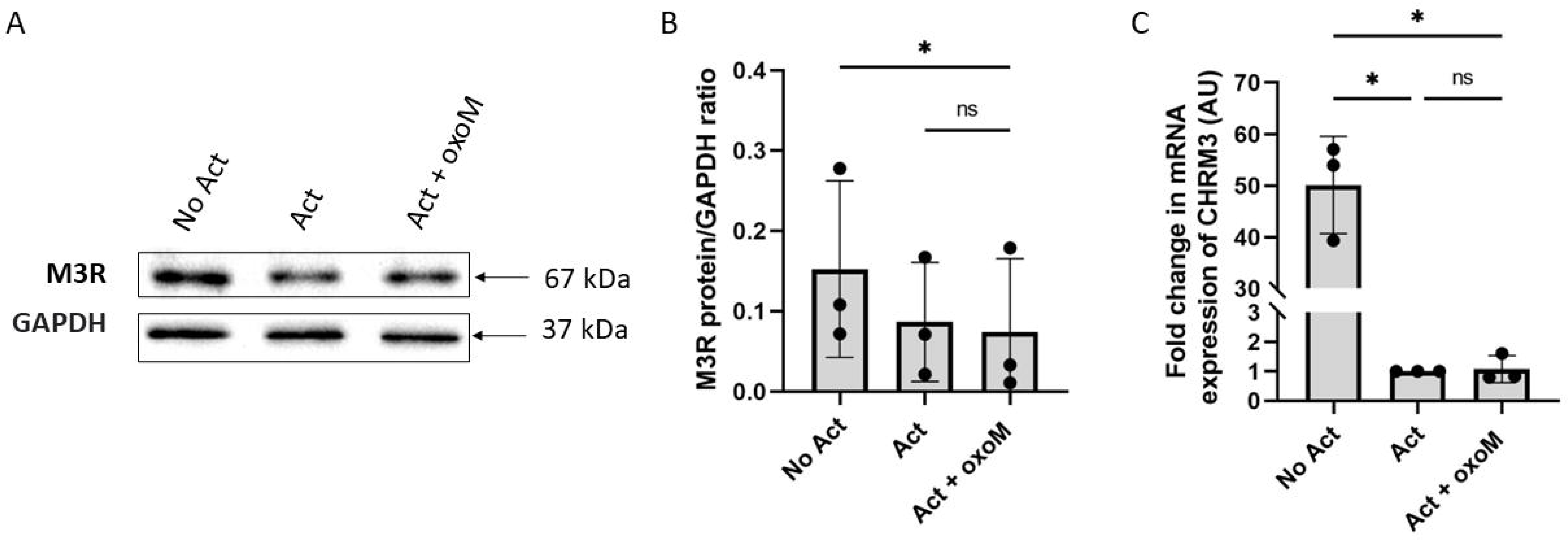
The effect of muscarinic receptor stimulation on M3R expression from memory Th cells. (A) Immunoblotting image of M3R protein from memory Th cells which not activated (No Act) or activated with ImmunoCult reagent (Act) and incubated with oxotremorine-M for 5 days. (B) Quantification of the bands for the M3R protein normalized to GAPDH. GAPDH used as a protein reference. The data was aggregated from 3 participants. (C) The mRNA level of *CHRM3* measured by RT-qPCR from memory Th cells which were not activated (No Act) or activated with ImmunoCult reagent (Act) and incubated with oxotremorine-M for 5 days. To determine the relative mRNA expression of M3R, the data was normalized to activated condition. Aggregated data normalized to activated condition is shown from memory Th cells of 3 participants. One-way ANOVA with Tukey’s multiple comparison test. *p< 0.05. ns, not significant.

**Figure 6.**
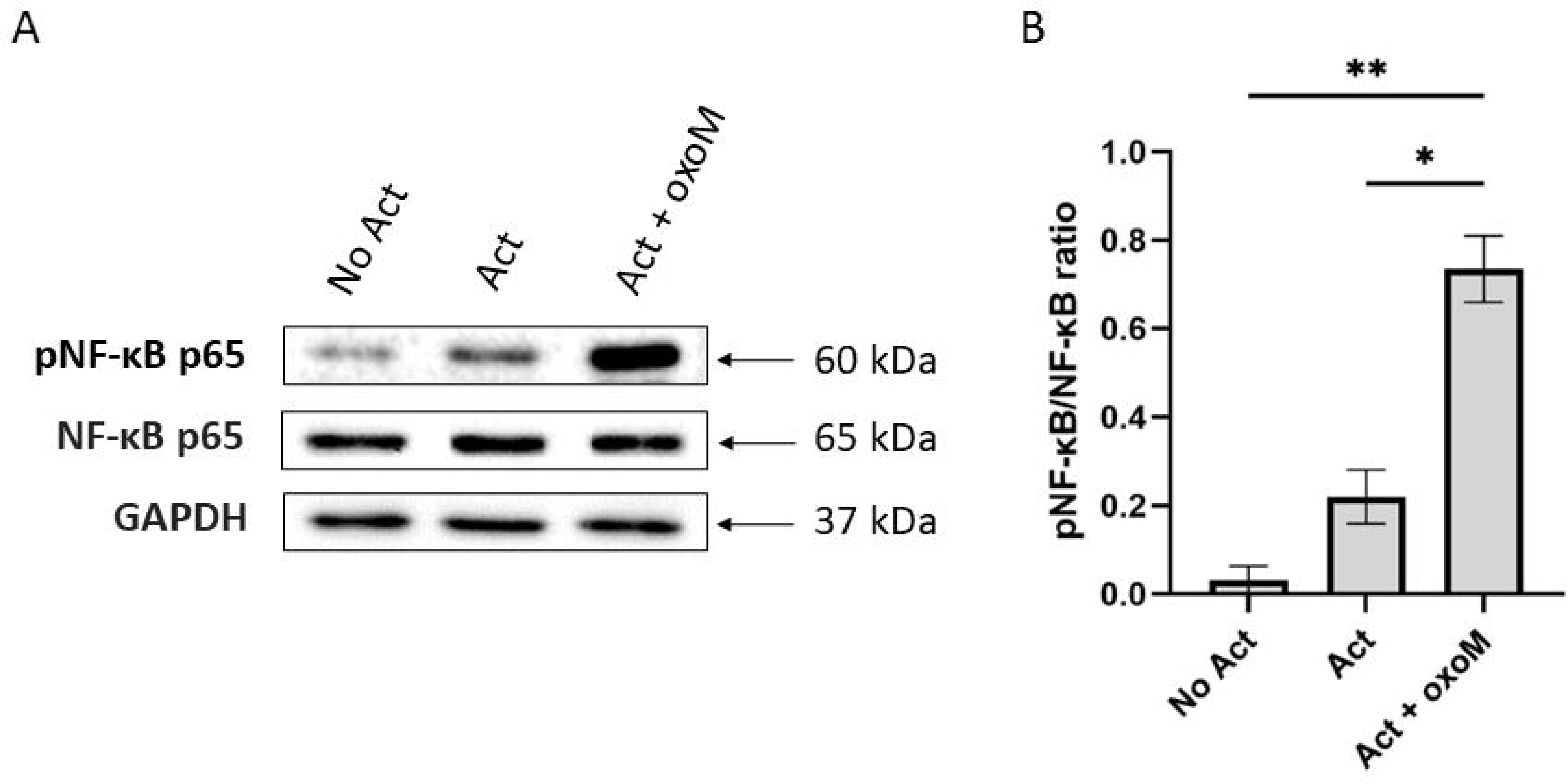
The effect of muscarinic receptor stimulation on phosphorylation of NF-κB p65 from memory Th cells. (A) Immunoblotting image of total and phosphorylated ser529 NF-κB p65 protein from memory Th cells which not activated (No Act) or activated with ImmunoCult reagent (Act) and incubated with oxotremorine-M for 15 mins. (B) Quantification of the protein levels of phosphorylated NF-κB (pNF-κB) p65 with respect to total NF-κB p65. A representative experiment from 4 participants is shown. One-way ANOVA with Tukey’s multiple comparison test. *p<0.05, **p<0.01.

### Effects of CHRM3-knockout on IFN-γ, IL-17A, and IL-4 secretion by memory Th cells

To directly demonstrate the role of M3R in memory Th cells, and to support the J10 blocking data, *CHRM3* was disrupted using CRISPR-Cas9 gene targeting by electroporation with RNP Cas9 complex. Oxotremorine-M failed to augment IFN-γ or IL-17A in the *CHRM3*-knockout condition as compared to control condition which indicates a direct involvement of M3R in the oxotremorine-M effect on memory Th cells (Figure 7 A, B). In contrast, oxotremorine-M continued to inhibit IL-4 in the *CHRM3*-knockout condition (Figure 7 C). Thus, the gene editing data supports a pro-inflammatory role for M3R with IFN-γ and IL-17A, while the role on IL-4 is M3R-independent. RT-qPCR analysis confirmed that CRISPR-Cas9 editing successfully reduced *CHRM3* expression to undetectable levels in the *CHRM3*-knockout cells compared to the control and nt-gRNA conditions (Figure 7 D). Cell viability remained remarkably constant throughout the days following electroporation (Figure 7 E). To further validate the CRISPR-Cas9 gene editing method, *TRAC* gene was targeted with specific gRNA or non targeting gRNA. *TRAC*-knockout efficiency was up to 90% in the *TRAC*-targeting group compared to the control and nt-gRNA groups based on flow cytometry analysis of TRAC protein expression (Figure 7 F, G).

**Figure 7.**
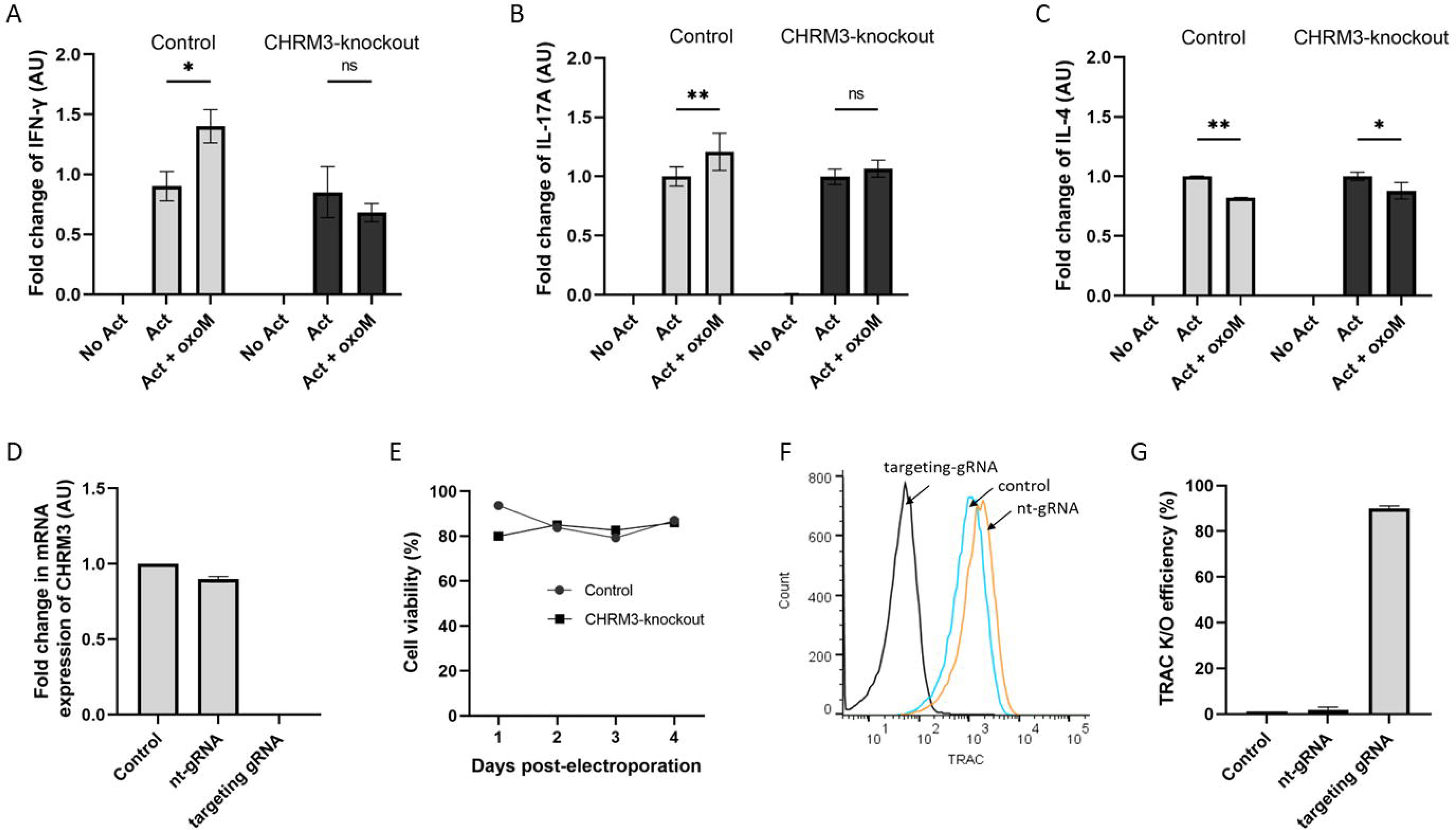
The effect of muscarinic receptor stimulation on *CHRM3*-knockout memory Th cells. *CHRM3* was knocked out genomically from primary memory Th cells using CRISPR-Cas9 gene targeting. Memory Th cells from control and *CHRM3*-knockout conditions were not activated (No Act), activated with ImmunoCult reagent (Act) and incubated with oxotremorine-M. The cytokine level of (A) IFN-γ, (B) IL-17A, and (C) IL-4 in the control and *CHRM3*-knockout conditions was measured by using ELISA. (D) Comparison of *CHRM3* mRNA expression from control, non-targeting gRNA (nt-gRNA), and *CHRM3*-knockout conditions. (E) Viability measurements over 4 days post-electroporation were recorded, comparing control and *CHRM3*-knockout. (F) Expression of TRAC protein in *TRAC*-knockout comparing control and nt-gRNA conditions with flow cytometry. (G) Compiled data of TRAC protein expression expressed as efficiency. A representative experiment from 2 participants is shown. Two-way ANOVA with Tukey’s multiple comparison test. *p<0.05, **p<0.01. ns, not significant. K/O = knockout.

## Discussion

Immune cells express muscarinic receptors but how the cholinergic system regulates their function is not completely understood. We addressed the expression and functional implications of muscarinic receptors, with a specific emphasis on the role of M3R in regulating the production of IFN-γ, IL-17A, and IL-4 from human memory Th cells. These cytokines were selected as prototypical markers representing the functional polarization of Th1, Th17, and Th2 cell subsets, respectively. Our findings revealed that stimulation of mAChRs led to an elevation in pro-inflammatory cytokines IFN-γ and IL-17A, dependent on M3R activity and involving NF-κB p65 activation. Conversely, mAChR stimulation suppressed IL-4 production in a manner sensitive to atropine but not dependent on M3R.

Memory Th cells were enriched by immunopanning from PBMCs for a phenotype of CD3^+^CD4^+^CD45RA^-^ CD45RO^+^. There are two main subsets of memory Th cells including central memory T cells which are CCR7^+^CD62L^+^CD45RA^-^CD45RO^+^ and effector memory T cells which are CCR7^-^CD62L^-^CD45RA^-^CD45RO^+^ (30). Since our study used an immuno-panning method, the phenotype of memory cells would include subtypes of CCR7^+/-^ and CD62L^+/-^ memory T cells. Memory Th cells were studied because they are thought to be one of the underlying causes of autoimmune diseases such as multiple sclerosis (MS) and psoriasis indicating their potential significance in autoimmune diseases (31–33).

To our knowledge, our data is the first description of mAChRs expression in primary human memory Th cells. A study on pan T cells from asthmatic patients showed that all five muscarinic receptors were expressed with M2R and M4R the highest (1). It is possible that the relative expression of mAChRs changes as Th cells transition from effector to memory. Our results showed a significant reduction in protein and gene expression level of M3R upon T cell activation. A similar study on naïve Th cells demonstrated that polarization is accompanied by a decrease in M3R expression (34). The consistent observation that T cells downregulate M3R suggests that endogenous ACh production might induce receptor desensitization.

Our data demonstrated a difference in muscarinic receptor usage between the cytokines. IFN-γ and IL-17A were M3R-dependent. Studies with murine systems showed that muscarine, an mAChR agonist, increased IFN-γ^+^CD4^+^ T cells in gut-associated lymphoid tissues, although it was not prevented by atropine (35). Muscarinic stimuli increased plasticity of Th cells towards Th17 polarization of naïve Th cells in mice (34). Thus, cholinergic signalling appears to have a consistent pro-inflammatory effect across human and mouse T cells. IL-4 secretion was M3R-independent although atropine-sensitive to oxotremorine-M. Atropine is a pan-specific muscarinic receptor antagonist with equal affinity for all subtypes of mAChRs. It is likely that one of the other receptors, M1R, M2R, M4R or M5R is responsible for the effects of oxotremorine-M on IL-4. The intracellular cytokine analysis of IL-4 showed atropine partially blocked the effect of oxotremorine-M suggesting that the block might be particular to cytokine secretion into the supernatant rather than affecting the proportion of IL-4-expressing memory Th cells.

J104129 is a potent M3R antagonist recognized for its highest affinity towards M3R (Ki= 4.2 nM) (19), has been used to block M3R and targeting M3R can improve symptoms in a multiple sclerosis model (36, 37), indicating the potential utility of M3R for the treatment of demyelinating diseases. Using a CRISPR-Cas9 gene targeting approach, we confirmed the M3R-blocker data with an effective knockout of *CHRM3* in the memory Th cells. These observations align with the findings of Nguyen *et al.*, who documented that M3R stimulation induces IFN-γ expression in CD4^+^ T cells in mice with increased NF-κB p65 activation (38). In Darby *et al.*, M3R was required for optimal immunity against helminth infection in a mouse model. However, the steady-state immune cell populations and *in vitro* Th2 polarization were normal in M3R^-/-^ mice. Furthermore, the authors showed that oxotremorine-M increased IL-4 in CD4^+^ T cells taken during helminth infection (22). Our results demonstrated that oxotremorine-M decreased IL-4 in an M3R-independent manner. However, there were significant differences as we studied healthy human participant samples treated *ex vivo* with a polyclonal activator. As evidence continues to emerge in this growing field of study, it appears that muscarinic receptor usage and functional outcome will depend on species, type of activation, the context of adaptive immunity, and experimental model.

In conclusion, our results demonstrate that muscarinic signaling increases the phosphorylation of NF-κB p65 and enhances the secretion of pro-inflammatory cytokines IFN-γ and IL-17A, with a corresponding decrease in IL-4 in memory Th cells. M3R is required for pro-inflammatory effects, while IL-4 decrease is atropine-sensitive. Understanding the role of the cholinergic system in memory Th cells allows for the development of new strategies to control immune responses, particularly in autoimmune diseases, by selectively blocking M3R.

